# A high throughput system reveals distinct segmentation clock phase responses in hiPSC-derived organoids

**DOI:** 10.64898/2026.07.21.739756

**Authors:** Rosie L Gallagher, Hedda A. Meijer, Adam Hetherington, Margarita Kalamara, Lindsay Davidson, Alistair Langlands, J Kim Dale, Philip J Murray

**Affiliations:** University of Dundee, Division of Molecular, Cell and Developmental Biology; University of Dundee, Mathematics; University of Edinburgh, LifeArc Centre for Rare Respiratory Diseases; University of Dundee, Human Pluripotent Stem Cell Facility; University of Dundee, National Phenotypic Screening Centre

## Abstract

During somitogenesis, the vertebrate body axis segments into transient periodic structures known as somites. Somite formation is regulated by a multicellular molecular oscillator known as the segmentation clock. Recent advances in human induced pluripotent stem cell (hiPSC) culture have shown that hiPSC-derived somitogenesis organoids (somitoids) exhibit segmentation clock oscillations and can be produced at scale, making them an excellent model system for high throughput investigation of the mechanisms underpinning the segmentation clock. However, somitoids interact both biochemically and mechanically, and accurate high-throughput sampling of segmentation clock phases is required to exploit this system effectively. Here we address these challenges using an image-based, high-content screening workflow. Individual hiPSC-derived somitoids carrying a segmentation clock reporter are cultured in 384-well plates, and a programmable feeding schedule is used to initiate oscillations that are monitored using fluorescence microscopy. We develop an automated pipeline for image segmentation and data analysis, represent oscillations using a compact set of parameters, and construct predictive mathematical models to interpret data. We find that: (i) a staggered feeding schedule that sequentially initiates oscillations yields large numbers of somitoids at defined stages of the segmentation clock cycle; (ii) media exchange in established oscillations induces a Type 0-like phase response, resetting the segmentation clock to a state characterised by low NOTCH pathway transcription; and (iii) control wells in media exchange experiments exhibit a Type 1 phase response in which the segmentation clock is delayed non-uniformly across the cycle. Using a mathematical model of segmentation clock dynamics along the anterior-posterior axis, we show that periodic activation of a Type 1 phase response could segment a continuous phase gradient — an insight with potential implications for the determination of somite boundaries *in vivo*.

**IMPORTANT:**

- Manuscripts submitted to Review Commons are peer reviewed in a journal-agnostic way.
- Upon transfer of the peer reviewed preprint to a journal, the referee reports will be available in full to the handling editor.
- The identity of the referees will NOT be communicated to the authors unless the reviewers choose to sign their report.
- The identity of the referee will be confidentially disclosed to any affiliate journals to which the manuscript is transferred.

**GUIDELINES:**

- For reviewers: https://www.reviewcommons.org/reviewers
- For authors: https://www.reviewcommons.org/authors

**CONTACT:** The Review Commons office can be contacted directly at

## Background

Somitogenesis is an important process that arises during the development of the vertebrate embryo. During somitogenesis, the presomitic mesoderm (PSM), bilateral strips of tissue that lie adjacent to the notochord and neural tube (Brent & Tabin, 2002; Chal & Pourquié, 2009; Christ & Scaal, 2008), segments at periodic time intervals to form transient structures called somites. These eventually differentiate into muscle, bone, and parts of the dermis. The rate of segmentation is species specific: approximately 30 minutes in zebrafish (Delaune et al., 2012), 90 minutes in chick (Palmeirim et al., 1997), 2 h in mice (Takashima et al., 2011), and 5 h in humans (Chu et al., 2019; Diaz-Cuadros et al., 2020; Sanaki-Matsumiya et al., 2022).

Somitogenesis is regulated by multiple signalling pathways. Opposing gradients of FGF/Wnt and RA pathway components extend along the PSM and play an important role in establishing the anterior-posterior (AP) axis (Pourquié, 2022). The segmentation clock, a molecular oscillator characterised by oscillating gene expression in the Fgf, Notch and Wnt signalling pathways, regulates the length of formed segments. In numerous species, it has been shown that expression of Notch/Fgf target genes (e.g. *HES7, NRARP, LFNG*) occurs out-of-phase with Wnt target genes (e.g. *AXIN2, DKK1*, Dequéant et al., 2006; Sewell et al., 2009).

Somitogenesis was first characterised in animal model systems (e.g. mouse, chicken, and zebrafish embryos (see Pourquié, 2022 for review)). In these models, assays often involved techniques such as *in situ* hybridisation and immunocytochemistry to characterise spatial patterns of clock gene expression at the mRNA or protein level (e.g. Takada et al., 1994; Aulehla et al., 2003; Bessho et al., 2001; Bone et al., 2014). Techniques such as microarrays have been used to globally characterise segmentation clock gene expression, (e.g. Dequéant et al., 2006). Hence, many molecular pathways that comprise the segmentation clock have been identified.

Advances in the development of human induced pluripotent stem cell (hiPSC) lines with segmentation clock reporters (e.g. HES7-Achilles, Venus-MSGN (Diaz-Cuadros et al., 2020) and HES7-luciferase (Matsuda et al., 2020)) have enabled the characterisation of segmentation clock oscillations in cell culture systems (e.g. Diaz-Cuadros et al., 2020; Matsuda et al., 2020). These protocols enable the generation of vastly increased amounts of biological material compared with *in vivo* studies. Hence, experimental techniques (e.g. OMICs approaches) that require relatively large amounts of biological material can be applied.

Somitogenesis organoids have allowed for many aspects of human somitogenesis to be studied (Meijer et al., 2025; Miao et al., 2023; Sanaki-Matsumiya et al., 2022; Yamanaka et al., 2023). Organoid models have been validated by characterising the major signalling pathways, including Wnt, FGF and Notch, and by phenotyping using techniques such as RNA-seq and time-lapse imaging. The measured segmentation clock period in human organoid models is approximately 5 h, a result that is consistent with cell-based studies (Diaz-Cuadros et al., 2020). Somitoids, as well as axioloids/segmentoids, can produce somite-like structures if cultured on extra-cellular matrix compounds such as Matrigel (Meijer et al., 2025; Sanaki-Matsumiya et al., 2022). Hence, somitogenesis organoids can recapitulate many important aspects of *in vivo* behaviour.

Despite the opportunities offered by somitoid model systems, the physical and biochemical interactions between co-cultured somitoids represent potential confounding variables that restrict high-throughput quantification. When micropatterned substrates have been used to constrain somitoids to sit in particular geometric patterns, it has been shown that signalling gradients are reinforced by neighbouring somitoids (Yaman & Ramanathan, 2023). These data suggest that neighbouring somitoids interact biochemically, presumably via the secretion of diffusible molecules. Moreover, in our hands, physically adjacent somitoids often merge.

High content screening (HCS) is an experimental design approach that combines high-throughput assays with automated microscopy in order to collect quantitative data from complex biological systems (Zanella et al., 2010). Traditionally this method has often been applied to drug discovery, where, typically, the response of cells/tissue to pharmaceutical treatments are assayed (Boutros et al., 2015). HCS architectures have been previously used to study organoid development (e.g., Sanaki-Matsumiya et al., 2022; Miao et al., 2023; Yaman & Ramanathan, 2023; Yamanaka et al., 2023). Whilst the larger sample sizes in high throughput studies offer possibilities for greater statistical robustness, post processing challenges include the need for automated data-handling and image analysis protocols at scale.

Segmentation clock oscillations in organoids can be initiated by environmental stimuli. Following media exchange, levels of HES7 expression have been found to increase over a short transient period, prior to the emergence of sustained large amplitude oscillations (Sanaki-Matsumiya et al., 2022). Delaying media treatment by 24 h resulted in a concomitant delay in the initiation of HES7 oscillations. Together, these data suggest that the controlled initialisation of oscillations via media treatment offers a means of generating high-throughput samples at different stages of the segmentation clock cycle.

The circadian clock literature provides tools in which to characterise the response of oscillators to perturbation (Winfree, 1980). Phase transition curves (PTCs) plot the phase of an oscillator at the time of perturbation against the phase after the perturbation. Closely related phase response curves (PRCs) describe how the oscillator phase responds to a stimulus applied at a given phase. A Type 0 phase response is characterised by a *hard reset* of the oscillator to some point on the cycle. In contrast, Type 1 PRCs are characterised by small phase perturbations of a constant sign at different points in the oscillatory cycle (i.e. either phase delays or accelerations but not both). Using mouse explants, phase response curves have been inferred from experiments in which cells from different PSM explants are merged or externally perturbed using microfluidics (Ho et al., 2024; Sanchez et al., 2022).

The availability of sufficient quantities of tissue has limited the application of high throughput omics techniques to the study of many developmental processes. However, iPSC-derived organoid models can allow for this barrier to be overcome. The aims of this study are to: (i) develop a high-throughput system that will enable automated analysis of isolated somitoids in standardised micro-environments; (ii) stage somitoids across the segmentation clock cycle; (iii) develop an automated data analysis pipeline that allows the human segmentation clock to be represented by a small number of parameters; and (iv) develop a mathematical framework that allows for characterisation of response of segmentation clock to external stimuli.

## Results

To generate high throughput quantities of human PSM tissue, hiPS cells with a HES7-Achilles reporter (Diaz-Cuadros et al. (2020)) were plated in an inclined 384 well plate (Day −3) to allow for the formation of embryonic bodies (EBs), followed by differentiation on Day 0. After media change, live imaging commenced on Day 2, during which the HES7-Achilles signal was detected using fluorescence microscopy (see Materials and Methods and Supplementary Figure S1 for a representative view of plate). Upon computing the mean HES7-Achilles signal in each well, sustained oscillations were observed with a period of approximately 4.5 h (data not shown), in agreement with previous observations (Meijer et al., 2025).

To improve the signal-to-noise ratio, the Cellpose segmentation algorithm (Stringer et al., 2021) was trained to identify and track somitoids (Figure 1 A and Materials and Methods). The mean HES7-Achilles signal in the somitoid domain was computed at each imaging time step (Figure 1 B1). After detrending (Figure 1 B2), a wavelet analysis (see Materials and Methods) was performed and the resultant time series was well fitted by a wavelet reconstruction (*R*^2^ = 0.82, Figure 1 B3). The fitted period is approximately 4.5 h, and the oscillator phase is estimated using a ridge detection method (Figure 1 B4). Hence, we implemented an automated pipeline that parameterises key features of segmentation clock oscillations in a high-throughput, hiPSC-derived somitoid model.

**Figure 1.**
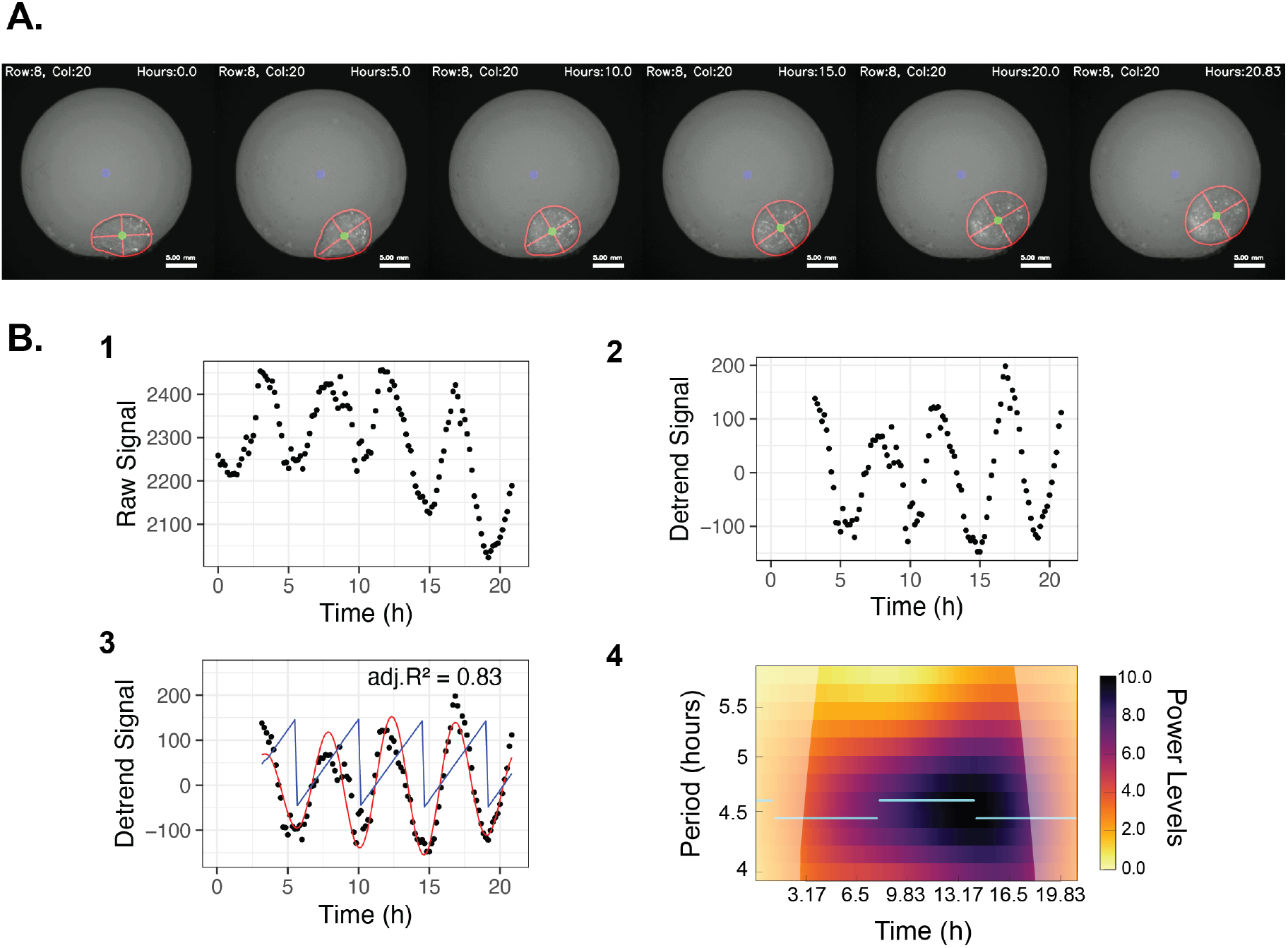
Signal processing and statistical modelling of HES7-Achilles signal in a single somitoid. **A**. Snapshots of a segmented somitoid (see Supplementary Movie S1) imaged on the YFP channel (used to capture HES7-Achilles) at 5 h time intervals. Somitoid boundary (red contour) plotted together with major and minor axes (red lines). Blue dot - centre of the well, green dot - somitoid centroid. **B**. Time series representing processing of the HES7-Achilles signal. **B.1** Mean raw signal. **B.2** Detrended signal. **B.3** Detrended signal with wavelet reconstruction. **B.4** Periodogram of the wavelet transform. Cyan lines represent the ridge where signal power is maximal.

To test the hypothesis that synchronous feeding yields synchronised segmentation clock oscillation, somitoids in a 384 well plate underwent simultaneous media exchange followed by 16 h of live-imaging (Figure 2 A). We found that the segmentation clock phase was approximately homogeneous across the multi-well plate (Figure 2 B). Upon computing the average phase in each column of the plate, we observed clear, synchronous oscillations (see the horizontal bands in Figure 2 C), indicating that HES7-Achilles is expressed simultaneously across the plate. These observations are consistent with previous reports of media change initiating segmentation clock oscillations (Sanaki-Matsumiya et al., 2022).

**Figure 2.**
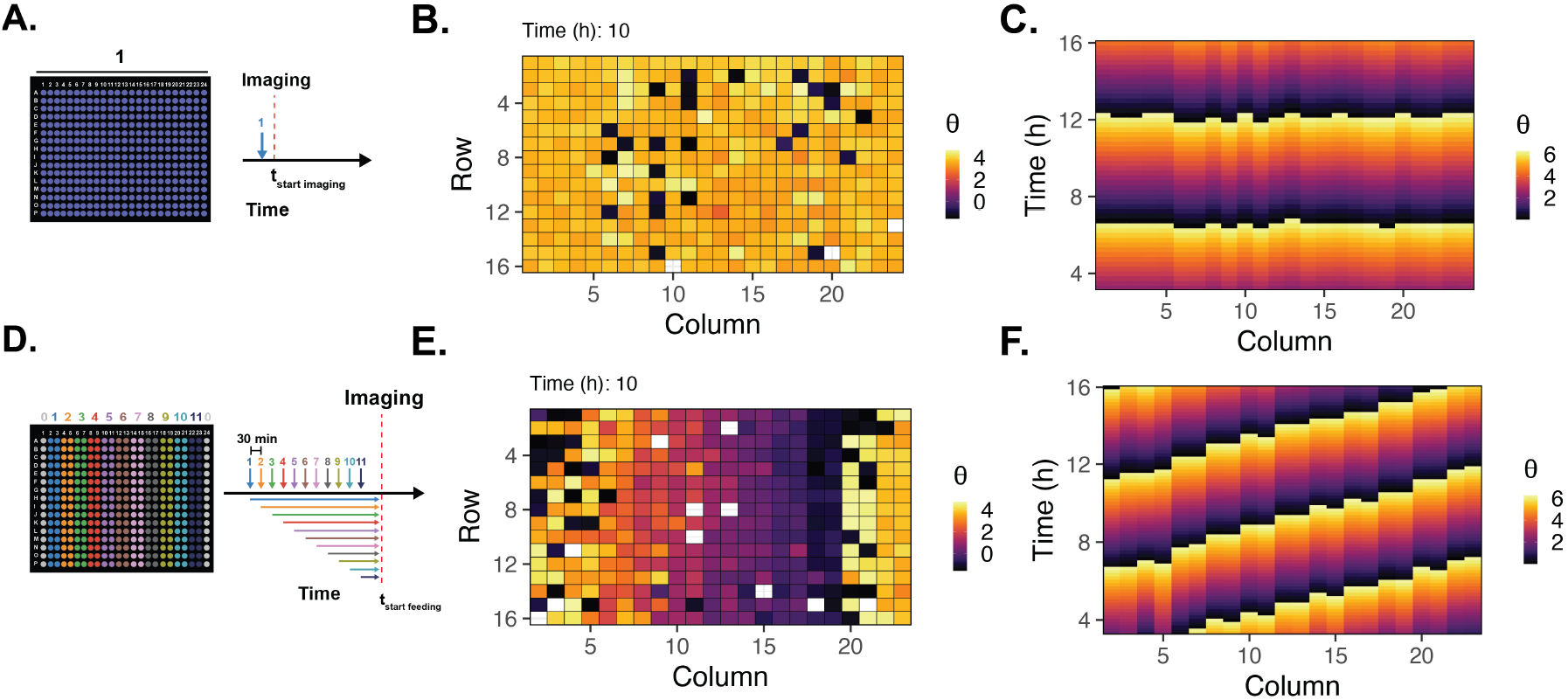
Staggered feeding induces phase shifted oscillations across 384 well plate. **A**. Schematic diagram of the simultaneous media change experiment. **B**. A 384 well grid view of the phase, *θ*, sampled at 10h (see Supplementary Movie S2 for time series). **C**. Phase averaged in each column is plotted against time. **D**. Schematic diagram of the 30 min staggered media change experiment Columns 1 and 24 are no feed control wells (grey). **E**. A 384 well grid view of the phase, *θ*, sampled at 10h (see Supplementary Movie S3 for whole time series). **F**. Phase averaged in each column is plotted against time.

To test if staggering media change results in a multi-well plate with somitoids that are distributed across the segmentation clock cycle, somitoids were assigned to feeding groups that were initiated at 30 minute intervals using a liquid handling robot (see adjacent pairs of columns in Figure 2 D) followed by live imaging. We identified a phase delay that increased approximately linearly across columns of the plate (i.e. later fed groups were delayed, Figure 2 E). In contrast to Figure 2 C, after averaging over rows, we observed a diagonal banding pattern in HES7-Achilles expression, consistent with a phase delay in later-fed groups (Figure 2 F). Note that Columns 1 and 24 acted as no feed controls and are omitted from the image. These results show that fine-grained temporal control of initiation yielded a multi-well plate with somitoids that are staggered across the segmentation clock cycle.

To validate the staggered initiation data, an additional set of experiments was performed in which live imaging was followed by qPCR in each feeding group (the plate was harvested 20 min after the last imaging time point, see Figure 3 A). We found that peaks and troughs in expression patterns were consistent with the NOTCH (*NRARP* and *HES7*) and WNT (*DKK1*) signalling pathways oscillating out-of-phase (Figure 3 B, (Matsuda et al., 2020)). Moreover, by extrapolating the phase measured at the end of live imaging to the time at which PCR was performed (i.e. 30 min later), the PCR and HES7-Achilles data were systematically aligned (Figure 3 B). As expected, peaks of HES7-Achilles (protein reporter) levels followed the peak in *HES7* mRNA with a delay of about a quarter of a cycle (i.e. approx. 1 h, Figure 3 C). These data confirm that staggered initiation yields a phase shift in multiple known molecular components of the segmentation clock and in distinct molecular pathways.

**Figure 3.**
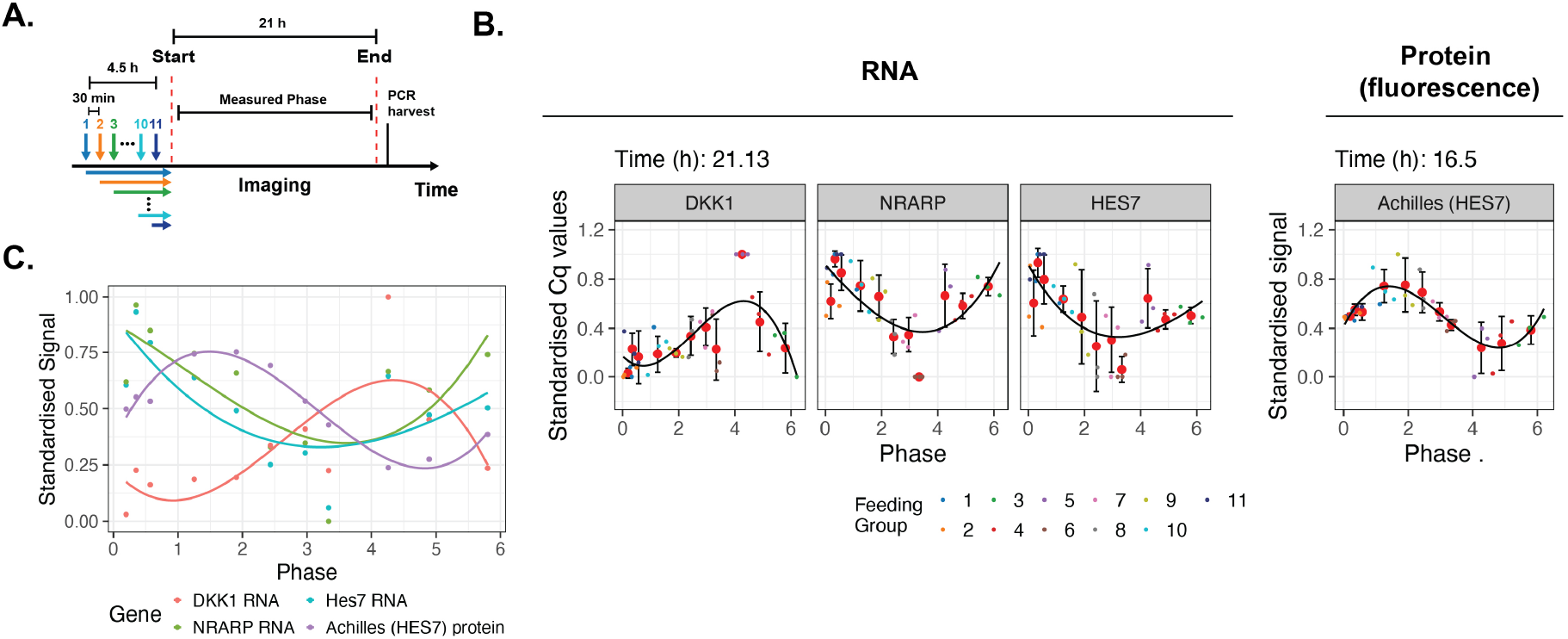
qPCR validation of somitoid oscillations after staggered feeding. **A**. A schematic illustration of the schedule. **B**. Left - expression of selected clock genes (DKK1 (Wnt), NRARP and HES7 (Notch)) quantified using qPCR is plotted against phase. Right - HES7-Achilles (protein) levels plotted against phase. Phase is extrapolated to qPCR sampling time. **C**. Overlay of data in B. Mean of standardised signal plotted against phase. Solid lines represent fitted sinusoids.

We expected to find that uniformly spaced feeding times would yield somitoids that were uniformly spaced across the segmentation clock cycle (e.g. a 30 minute delay in initiation would delay the segmentation clock by the same amount). To test this hypothesis, the mean phase, 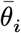, was computed for each feeding group 10 h post-imaging (Figure 4 A). We then quantified how the measured phase difference between consecutive feeding groups differed from the value expected given the feeding time step, *dt*_*f*_, i.e. the *phase deviation*

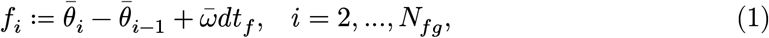

where 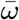 is the mean frequency. Under the null hypothesis that a feeding time delay of *dt*_*f*_ induces a phase delay of *ωdt*_*f*_, we expected to find that *f*_*i*_ = 0 for all *i*. After repeating the staggered initiation experiment in Figure 2, it was found that the staggered phase distributions were largely reproducible across experiments (see Figure 4 B for polar plot representations). However, upon computing the phase deviation curve (Equation 1) we observed a systematic bias, with a trough and peak at feeding times of approximately 1.5 and 3 h, respectively (Figure 4 C). These features of the phase deviation curve were reproduced (see Supplementary Figure S2) when: (i) the feeding time interval was varied (*dt*_*f*_ = 45, 50, 55 min); (ii) the assignment of feeding groups to columns was randomised; and (iii) the plate was horizontal from the time of differentiation onwards. The identified phase deviation was therefore a robust feature of the system.

**Figure 4.**
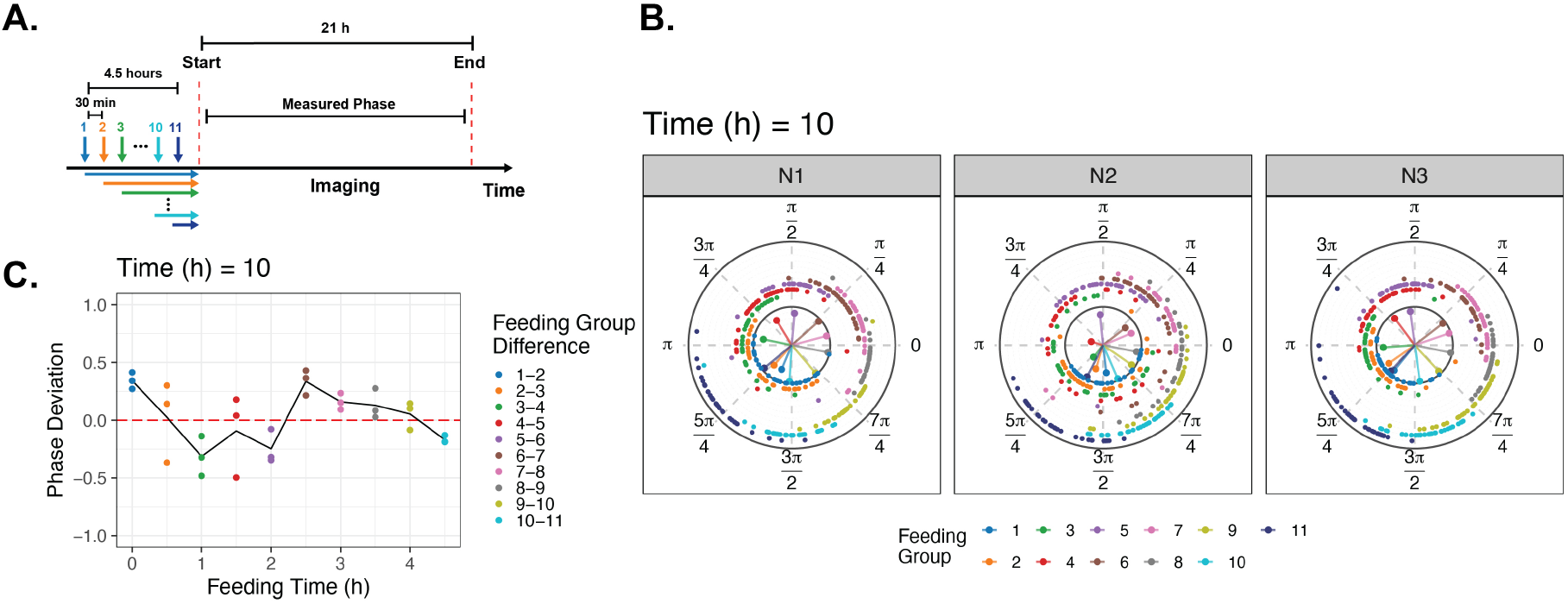
Uniformly staggered initiation yields non-uniform phase separation. **A**. A schematic illustration of the feeding schedule. **B**. Polar representation of segmentation clock phase measured at 10 h in 3 biological repeats (approx. 384 somitoid per repeat). Each datapoint in the outer spiral represents an individual somitoid. Radius scales with feeding group index for visualisation purposes only. Solid radial lines in inner circle represent the mean phase and order parameter within each feeding group. **C**. The phase deviation is plotted against feeding time (N=3, Equation 1). Solid line represent mean of experimental measurements.

To investigate the origin of the non-uniform phase deviation, the experimental design was modified to enable imaging before and after perturbations. The staggered initiation protocol described in Figure 4 was again used to obtain a multi-well plate with somitoids distributed across the segmentation clock cycle (Figure 5 A). Following live imaging for 21h, somitoids were assigned to one of two groups: media exchange or control (Figure 5 A). The multi-well plate was then moved from the imaging microscope to an adjacent robotic feeding station and incubated for 30 min. The selected wells (Rows 2,3,6,7,…) underwent media exchange (Figure 5 B). Subsequently, the plate was returned to the microscope and live imaged for a further 21h. After quantifying the segmentation clock phase, as described in Figure 1, we characterised the phase response to perturbation by computing phase transition distributions (PTD, i.e. joint probability distributions for the *phase after* given the *phase before*, see Materials and Methods) for both treatment groups.

**Figure 5.**
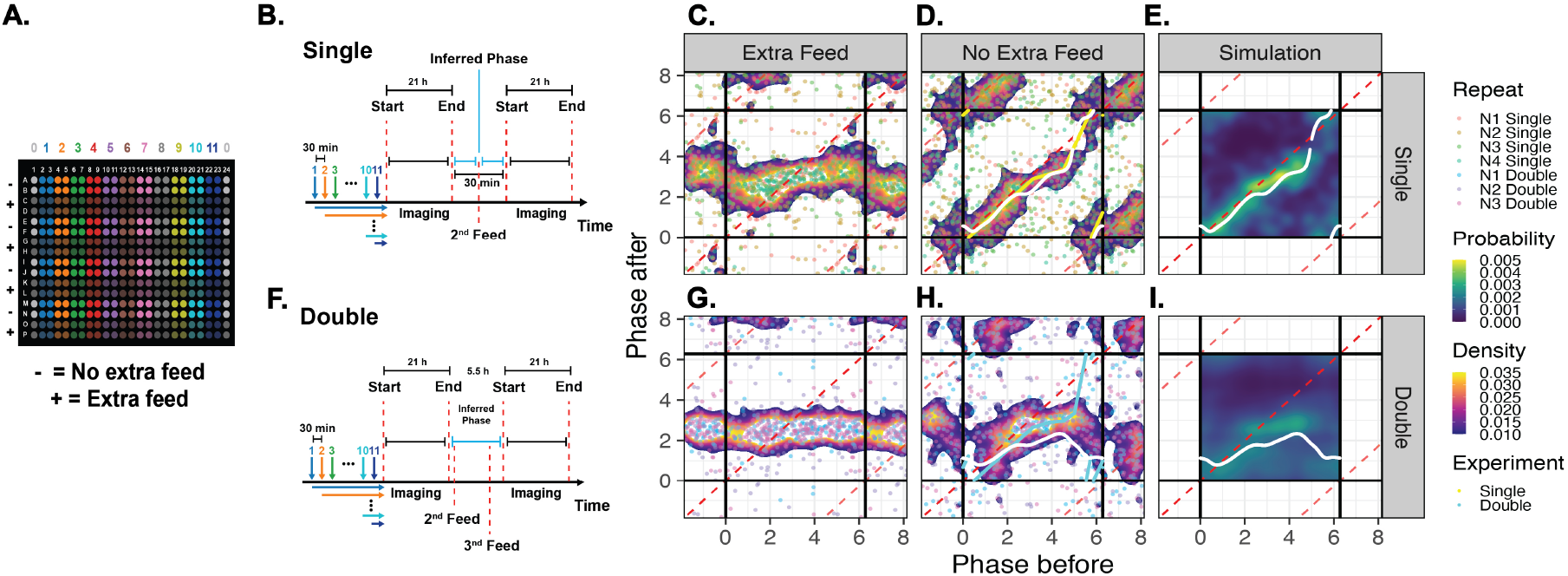
Phase response upon perturbing established oscillation. **A**. Plate layout and feeding group assignment. Signs indicate rows where extra feed takes place. **B-E**. Single extra feed. **F**,**I** Double extra feed. **B**,**F** Timeline of the feeding schedules. **C**,**G** Phase transition distributions (PTDs) for media change experiments. Dots represent extrapolated phase before and after perturbation. Heat map represents somitoid density. **D**,**H** PTDs for no-media change experiments **E**,**I**. Kernel density estimates for PTDs. Red dashed lines - no effect. White lines - expectation of PTD (Equation 8). Cyan lines - a fitted piecewise linear model of the PTC (Equation 12). Yellow line: predicted composite PTC, *Q* (Equation 15). See Supplementary Figures S3-S6 for biological repeats.

We found that in the media-exchanged wells the phase was reset to approximately 3 radians after media exchange (Figure 5 C, Supplementary Figures S3 and S4 (top row)). Although there was weak dependence on phase before, this behaviour resembles a Type 0 phase response. Aligning with the qPCR profiles in Figure 3, the reset point occurs at a phase associated with low levels of Notch transcriptional activity and increasing levels of Wnt transcriptional targets. Whilst it has previously been reported that media exchange initiates segmentation clock oscillations (Sanaki-Matsumiya et al., 2022), these data suggest that media exchange can reset the segmentation clock of established oscillations to a prescribed state.

For somitoids in the control wells, which did not experience media exchange, the PTD analysis reveals a small phase delay in the second part of the cycle (i.e. for *π* < *θ* < 2*π*, see Figure 5 ans Supplementary Figures S3 and S4). The fitted phase transition curve (PTC, *P*) was well-approximated by a piecewise linear model (Figure 5 D and Equation 12). The form of the PTC suggests that, independent of media exchange, the segmentation clock undergoes a bilinear delay in the second part of the cycle. In a perfect negative control, *P* (⋅) would be an identity transformation (i.e. the phase after equals the phase before so *P* (*x*) = *x*, see red dashed line in Figure 5 D and Supplementary Figures S5 and S6). Therefore, physical perturbation of the multi-well plate induces a non-uniform delay in the segmentation clock in a manner that is independent of media exchange.

To test the phase response model, we used the PTD in Figure 5 D to compute a composite PTD that predicts the phase response of two consecutive feeding perturbations (see composite PTD in Figure 5 I and compare with single in Figure 5 E, Materials and Methods). Notably, segments of the single PTC with slope less than one are predicted to be shallower in the composite PTC and *vice versa*. To test this prediction, a further experiment was performed in which the media exchange protocol at 21 h was repeated at 25.5 h post initiation of imaging (see Figure 5 F, Supplementary Figurs S5 and S6). We found that the outcomes of the double feed experiments were well described by the predicted composite PTCs (see Figure 5 G and H, Supplementary Figure S5 and S6) for both Type 0 and 1 phase responses. This analysis provides a validation of the phase response framework and suggests that the segmentation clock is competent to respond to a second stimulus within one clock cycle.

We next investigated if the identified PTC can explain the phase deviation curves presented in Figure 4 C. We hypothesised that a single stimulus applied at the end of the staggered feeding induces a Type 1 phase response; as the somitoids are in different phases of the cycle at the time the stimulus is applied, a differential response is observed (see Figure 6 A). To formalise this argument, the mathematical model was generalised to account for the phase increase experienced by each feeding group prior to the phase response. Let *θ*_*i*_(*t*) represent the phase of the *i*^*th*^ feeding group. Initialising the phase at *θ*_0_, just before the perturbation

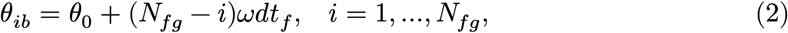

where *dt*_*f*_ is the feeding time interval and *ω* is the clock frequency. After the perturbation, an instantaneous phase update is given by

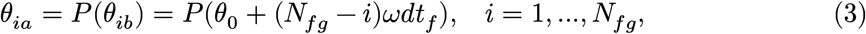

with *P* (.) as defined in Figure 5 D. When this nonlinear difference equation is solved graphically for different feeding groups (see Figure 6 B), one sees how non-uniform phase deviation can arise. Moreover, the computed solution can be represented via polar plot or a phase deviation curve (Figure 6 C and D, respectively) to enable comparison with the experimental data presented in Figure 4 C and D. Finally, after computing phase deviation curves for the different feeding times, we compared simulated phase deviation curves with the experimental measurements (see Figure 6 E). We can see that the simulated response curves provide a quantitative explanation for phase deviations observed in Figure 4. This analysis provides supporting evidence that a single PTC underlies the non-uniform phase responses observed in established and recently initiated segmentation clock oscillations.

**Figure 6.**
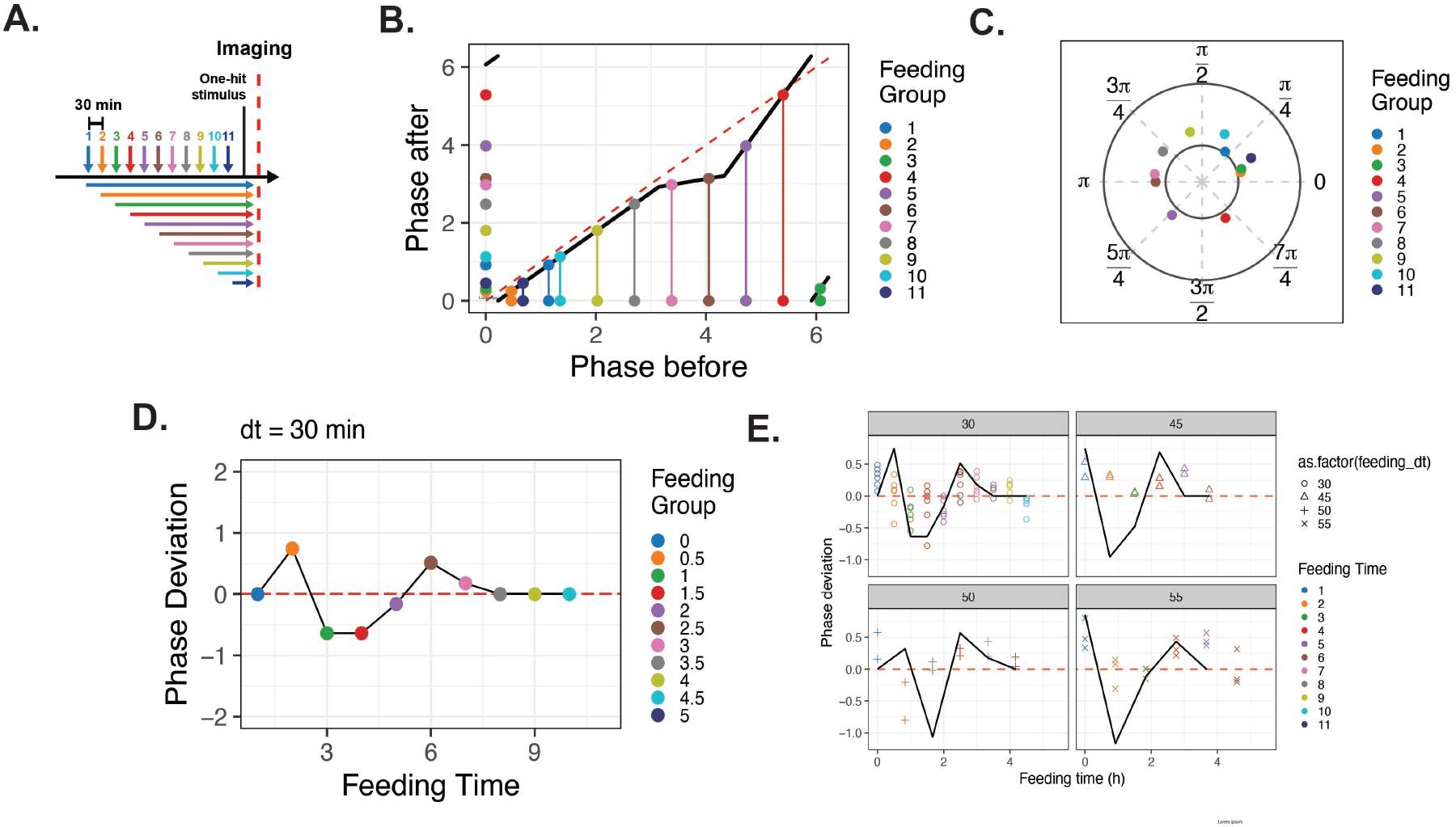
Modelling of a single Type 1 phase response post initiation explains phase deviations observed at different feeding time intervals. **A** Schematic illustration of experimental design. **B** Simulation of PTC applied to feeding groups that have uniformly sampled phases (Equation 3 with PTC given by Equation 12). Phase after perturbation is plotted against phase before. **C**,**D** Polar and phase deviation representations of the solution computed in B. **E** Comparison of simulated and experimentally measured phase deviation curves for different feeding time intervals. Experimental data (Figure 4 and Supplementary Fig S5). Simulations - solid black lines. Dashed red line: theoretical negative control.

Could the identified Type 1 phase response play an instructive role during embryo development? Given that the segmentation clock is competent to fully respond to a second stimulus applied one cycle after the first, we hypothesised that periodic activation of a Type 1 phase response could mediate a mechano-chemical feedback loop. To investigate this question, we first considered an *in silico* base-line clock and wavefront model of segmentation clock phase dynamics along the anterior-posterior axis in which a prescribed frequency gradient propagates posteriorly at a constant speed, *v* (see Figure 7 A and Materials and Methods). This model exhibits, as expected, anterior-propagating phase waves and a smooth, posteriorly propagating phase gradient (see Figure 7 B and C). When a periodic Type 1 phase response is imposed (see Figure 7 D and E), the model solutions show a *segmented* phase gradient (see step-wise pattern in Figure 7 F). In this model the periodic stimulus delays and entrains the segmentation clock to a longer period via repeated application of the Type 1 phase response. Hence a periodic Type 1 phase response applied in the context of a travelling phase gradient yields segmentation of a smooth phase gradient.

**Figure 7.**
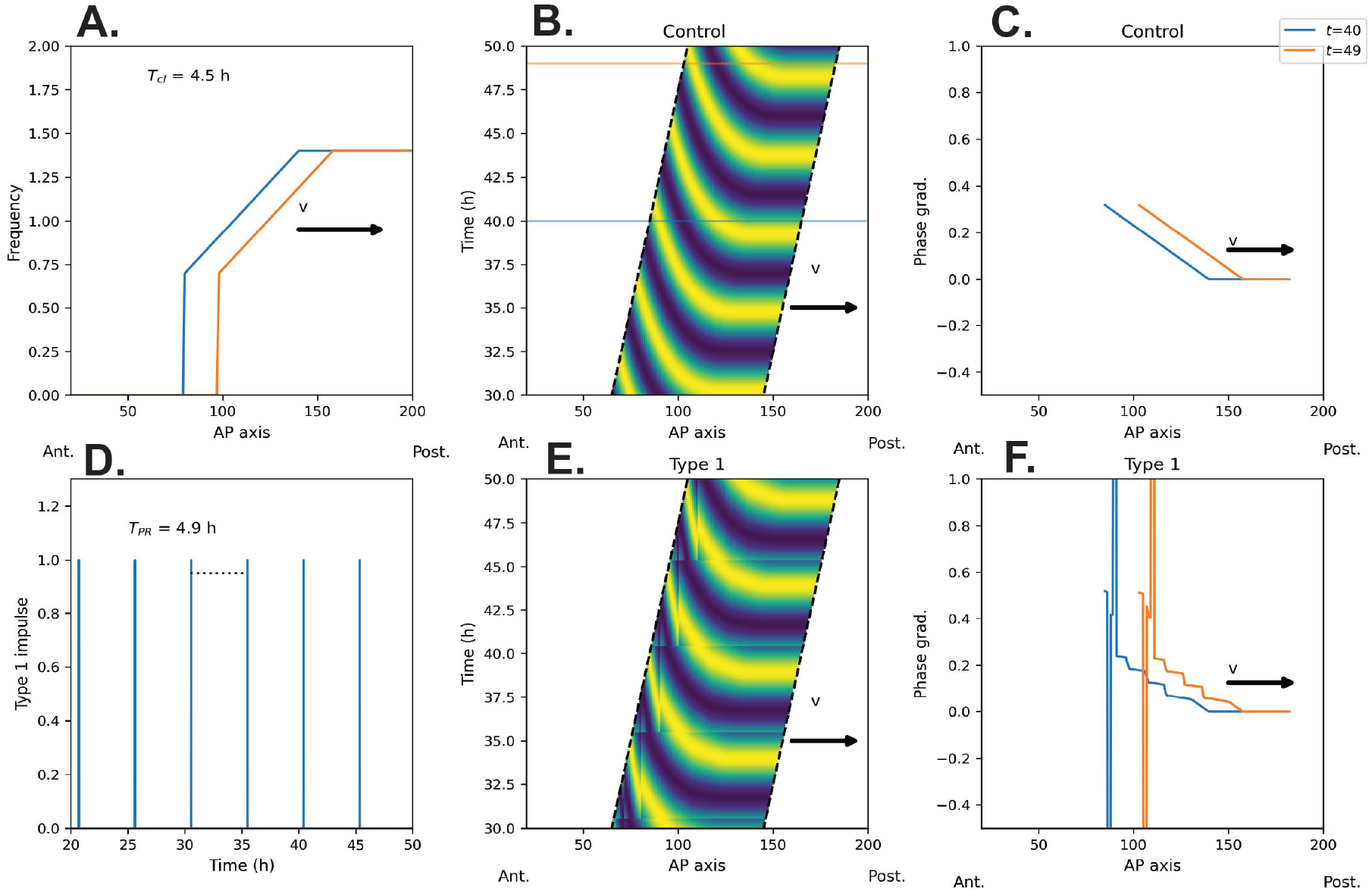
Periodic activation of Type 1 PRC yields a segmented phase gradient in a model of AP axis patterning. **A** Snapshots of frequency gradient travelling posteriorly at constant speed, *v*. **B** Oscillator phase dynamics plotted against AP axis position and time. **C** Snapshots of phase gradient plotted against AP axis position. **D** The Type 1 phase response is applied at periodic time intervals. **E** Oscillator phase dynamics plotted against AP axis position and time. **F** Snapshots of phase gradient plotted against AP axis position. Dashed lines denote anterior (left) and posterior (right) boundaries. See Table 1 for parameter values.

## Discussion

Somitogenesis is a fundamental developmental patterning process in which morphology is directly coupled to a complex molecular network that regulates the specification of a population of neuromesodermal progenitor cells. Whilst studies in model organisms (e.g. mouse, chicken, zebrafish (Delaune et al., 2012; Palmeirim et al., 1997; Takashima et al., 2011)) have led to the discovery of crucial regulating principles and the development of novel tools, the recent advent of hiPSC-derived somitogenesis organoids has allowed human somitogenesis to be studied at scale.

**Table 1.** A table with parameter values for the AP axis model.

| Parameter | Description | Value | Unit |
| --- | --- | --- | --- |
| $v$ | Frequency gradient velocity | 2 | c.d./h |
| $x_0$ | Frequency gradient offset | 0.0 | c.d |
| $L$ | Frequency gradient length | 60.0 | c.d. |
| $\omega_0$ | Seg. clock freq. | 1.4 | rad h<sup>-1</sup> |
| $T$ | Stimulus period | 4.9 | h |
| $m_1$ | PTC slope | 0.24 | Dimensionless |
| $m_2$ | PTC slope | 1.94 | Dimensionless |
| $c$ | PTC intercept | 6.05 | rad |
| $x_1$ | PTC break | 4.33 | rad |

In this study we have adapted methods from high content screening studies to enable a high-throughput characterisation of the somitogenesis clock. A robotic feeding system was used to initiate oscillations via media changes and an automated time-lapse imaging system was used to assay segmentation clock oscillations in somitoids grown in 384-well plates. Given that the overall size of the imaging dataset is around 250 GB per experiment and that we have analysed time series from thousands of somitoids, a bespoke automated image processing and analysis pipeline was developed for image segmentation, signal analysis, and interpretation.

Using the developed approach it has been shown that the initiation of the segmentation clock in hiPSC-derived somitoids can be controlled to yield a somitoid population distributed across the segmentation clock cycle. When validated by qPCR, we found, as expected, that the WNT and NOTCH pathways were expressed out of phase. In addition, we have found that: (i) staggered feeding can be used to activate the segmentation clock such that a population of somitoids are systematically distributed across the segmentation clock cycle; (ii) established oscillations exhibit a Type 0-like phase response to media change in which they are reset to a phase of the segmentation clock characterised by low levels of NOTCH and increasing levels of WNT transcripts; (iii) perturbing the plate in order to feed selected wells results in a Type 1 phase response.

The mechanism underlying the Type 1 phase response is not clear. Plate movement could result in mechanical stimulus owing to movement of the somitoids. Notably, YAP signalling has been shown to regulate segmentation clock oscillations (Hubaud et al., 2017). However, movement of the plate could also result in media mixing, allowing somitoids to obtain fresh metabolites or oxygen, which could affect the oscillation profile, as metabolic activity, particularly glycolysis, has been shown to control segmentation clock genes such as Hes-7 and somite formation (Anderson et al., 2025; Matsuda et al., 2025; Miyazawa et al., 2025). Additionally, we have not uncoupled the Type 0 and Type 1 phase responses, as the media-treated wells also experience the no-treatment effect. These questions could be addressed using, for example, inhibitor treatments that block specific signalling pathways and/or by studying phase response in modified genetic backgrounds. However, this is outside the scope of the current work.

It is an open question whether the identified Type 0 and 1 phase responses are relevant *in vivo*. A potential model is that a physical stimulus, such as the formation of segment pairs, acts as a mechanical stimulus that regulates the segmentation clock in the PSM via the Type 1 phase response curve identified in this study. In such a model, the Type 1 phase response acting on a phase gradient results in phase gradient segmentation.

## Conclusion

We have implemented a pipeline for high-throughput characterisation of somitogenesis, enabling studies on human somitogenesis that require large amounts of input material. A staggered feeding programme resulted in the production of samples which represent different phases of the segmentation clock. Although we have identified that the segmentation clock is sensitive to perturbations of the plate, the approach could be adapted to characterise the system’s sensitivity to, for example, chemical perturbations (e.g., inhibitors of key signalling pathways). Moreover, the hiPSC-derived somitoid system offers the possibility of using gene editing to characterise phase responses across different genetic backgrounds. Altogether, this system provides novel opportunities for the analysis of the segmentation clock.

## Material and Methods

### Experimental

#### Cell line, hiPSC maintenance and somitoid culture

The entire study was performed using HES7-Achilles hiPSCs. These cells express the YFP variant ACHILLES at the end of the HES7 open reading frame separated by a self-cleaving T2A peptide (Diaz-Cuadros et al., 2020). The hiPSCs were maintained by the Human Pluripotent Stem Cell Facility at the University of Dundee (see Meijer et al., 2025). Somitoids were generated from a previous protocol (Meijer et al., 2025) with the following modifications: hiPS cells were seeded into Akura 384 Spheroid microplates (InSphero) using 300 cells per well. The plate was centrifuged at 500xg for 5 minutes and swirled for 1 min to compact the cells and incubated on a 30 degree stand at 37 °C, 5% CO2 for 3 overnight incubations.

The resulting embryonic bodies were washed 2x with 50 *μ* l pre-warmed DMEM-F12 (Gibco) using the Biotek 405 LS microplate washer (Agilent) and the Multidrop Combi reagent dispenser (Thermo Fisher Scientific). Subsequently, the DMEM-F12 was replaced with 50 *μ*l pre-warmed SIB+ (DMEM/F12 [Gibco] supplemented with 1× N2 [Gibco], 1× B27 [Gibco] supplements, 1× nonessential amino acids [NEAAs; Gibco], 1× glutamax [Gibco], 1× sodium pyruvate [Gibco], 1× penstrep [Gibco], 10 μM SB431542 (Sigma), 10 μM CHIR99021 (Tocris), 2 μM DMH-1 (Tocris), and 20 ng/mL bFGF (PeproTech). After all washes the plates were centrifuged at 500xg for 30 seconds. After 2 overnight incubations with the plate on a 30 degree stand in the incubator at 37°C and 5% CO2, the somitoids were fed with 50 μl pre-warmed SIB (SIB+ without SB431542, CHIR99021, DMH-1 or bFGF; unless stated otherwise in the legends).

If somitoids were fed together, the Biotek 405 LS and Multidrop Combi were used. If feeds were staggered over time, somitoids were fed using the Bravo automated liquid handling platform (Agilent). For some experiments there were additional feed(s) the next day (see legends). Depending on the experiment, the somitoid retention rate for analysis ranged from 70% to 95% (see Supplementary Information).

#### RNA purification and qPCR

The somitoids were washed using 50 *μ*l PBS prior to lysis in 15 *μ*l RLT buffer (part of the RNeasy Micro purification kit, Qiagen) using the Biotek 405 LS and the Multidrop Combi. After 10 minutes the lysates were pooled (32 wells per sample) using the Bravo. RNA was purified using the RNeasy Micro purification kit (Qiagen) according to the manufacturer’s protocol including DNaseI treatment. Quality control of the RNA and the RT-qPCR were performed as previously described (Meijer et al., 2025). qPCRs were performed with primers for the clock genes PPP1CA, HES7, NRARP and DKK1. For primer sequences for PPP1CA, HES7 and NRARP see (Meijer et al., 2025). Primer sequences for DKK1 are DKK1-F1: ATGCGTCACGCTATGTGCTG and DKK1-R1: CTGGAATACCCATCCAAGGTGC. All samples were analysed using the ddCq method and all Cq values for triplicate wells were within 0.5 Cq. The results for HES7, NRARP and DKK1 were normalised for PPP1CA. No RT and water controls were included and contributed to less than 0.8% of the relative amounts obtained with the cDNA samples. For all primers the primer efficiency was checked and found to be within 101-108% over at least 3 orders of magnitude.

#### Time-lapse imaging

Somitoids were imaged on the Operetta CLS High-Content Analysis System (Revvity) using a 10x objective. The atmospheric control was set to 37°*C*, 5% CO2. Time lapse series were performed using the YFP filter and all somitoids were imaged in a single plane at ten minute intervals. Images were acquired on a per-well basis, with an 8-minute time delay from well 1 to well 384 (the first acquisition to the last image acquisition) in a given imaging time step. To compensate for the time delay, timestamps were extracted from the microscope metadata file. The somitoids were imaged before and after time lapse imaging using brightfield to enable analysis of somitoid shape and quality. Further details are provided in the Supplementary Information.

### Image processing

Image segmentation was performed using Cellpose with a custom fine-tuned model built on the cyto3 model (Stringer et al., 2021) and the HES7-Achilles signal was averaged across somitoid masks at each time step. Time series were detrended and wavelet analysis was used to identify oscillator phase. Further details are provided in the Supplementary Information.

#### Phase extrapolation for qPCR analysis

Imaging data were extrapolated by sampling phase at a time point close to PCRs’s harvest time (21.14h-4.5h) and extrpolating. Next, the detrended imaging data and phase *θ*(*t*) were averaged (or circular averaged) by experiment and the feeding group. The signal was then standardised using a min-max method as qPCR data. Finally, the standard deviations were calculated for the given signal. Further details are provided in the Supplementary Information.

### Mathematical Models

#### AP axis model

Consider a discretised spatial domain, *x*, representing the AP axis. Let *θ*(*x, t*) represent the phase at given point on the AP axis at time *t*. An piecewise-linear, imposed frequency profile is given by

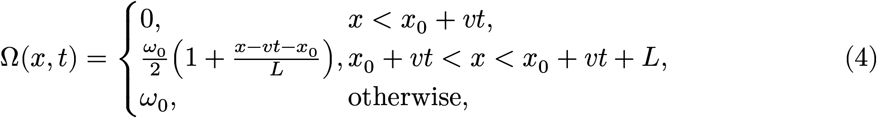

where *v* is frequency gradient velocity, *ω*_0_ is posterior frequency, *L* is the length of the frequency gradient and *x*_0_ is the position of the anterior boundary of the PSM at *t* = 0. The phase along the AP axis is obtained by integrating

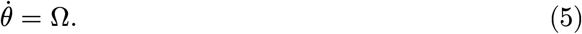

The initial condition is *θ*(*x*, 0) = 0. The Type 1 phase response fitted in (Figure 5 D, Equation 12) is imposed at *T* -periodic time intervals.

#### Computing summary statistics from the empirically estimated PTD

The segmentation clock phase time series before and after a perturbation are identified as described in Figure 5. The phases are linearly extrapolated to the perturbation time to define, for a given somitoid, the phase before, *θ*_*b*_, and the phase after, *θ*_*a*_. Upon measuring multiple somitoids, a population of phase before and after measurements are obtained 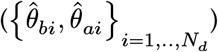. These data are used to empirically estimate the joint probability distribution, **P**(*θ*_*a*_ | *θ*_*b*_), from which further estimates are dervied.

To proceed, phase is discretised on a uniform domain

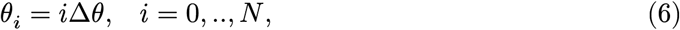

where Δ*θ* = 2*π*/*N*. A kernel density estimator is used to identify an empirical approximation to **P**(*θ*_*a*_ | *θ*_*b*_) in the form of a discrete phase transition matrix, *p*_*ij*_ (i.e. the probability of a somitoid transition from State *j* to State *i*).

Let *π*_*j*_ represent the probability of finding a somitoid in State *j*. After a single phase response, the probability of being in State *i* is

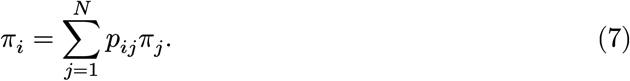

Given a somitoid is in State *j* before a perturbation, the expected phase after is

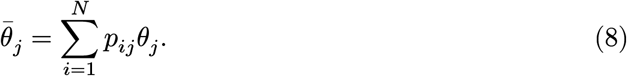

This form is analagous to a phase transition curve (PTC), i.e. a mapping from phase before to the expected phase after. This expectation is plotted in Figure 5.

Now consider application of a second perturbation, one cycle after the first. Under the assumption of 2*π* periodicity, the probability of a somitoid being in State *k* after consecutive perturbations is

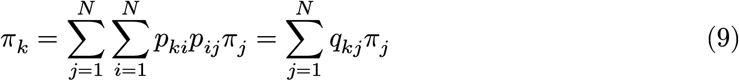

Here *q*_*kj*_ represents the matrix product

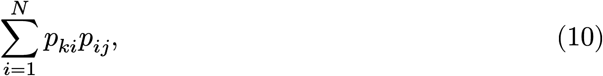

and the expectation for a composite feed is

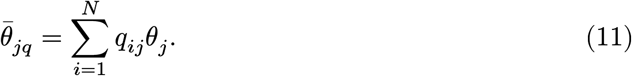

### A parametric form for the PTC

As an alternative to using the PTD, a parametric form has also been assumed for the PTC and fitted to the phase observations. A continuous piecewise linear PTC of the form

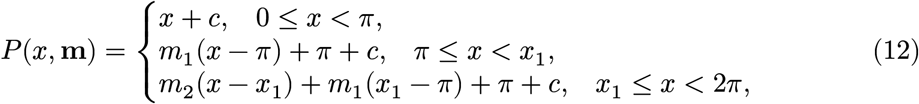

where **m** = (*m*_1_, *m*_2_, *x*_1_, *c*)^*T*^ .

There are two unknown slopes (*m*_1_ and *m*_2_), one break point (*x*_1_) and an intercept (*c*). To determine the unknown parameters, an objective function is defined to be

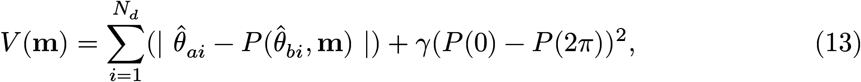

where the last term is a constraint that enforces continuity of *P* . The objective function is minimised using a Nelder-Mead algorithm implemented in R studio via the optimize package. Initial guesses are randomly sampled from uniform distribution of parameters and the *optimal* parameter set is chosen to be the one yields the overall minimum of *V* .

Letting *θ*_*b*_ and *θ*_*a*_ represent the phase of a somitoid before and after a perturbation, respectively, the instantaneous phase change induced by a perturbation is represented by the equation

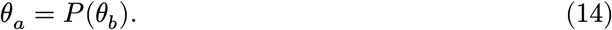

This equation is analagous to Equation 8.

Application of a second perturbation one cycle later yields a predicted *composite* phase response given by

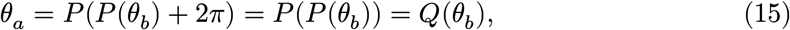

where *Q*(.) is a composite function that can be unambiguously computed given knowledge of P(.) (see Figure 5 H (cyan line) and I for predicted composite given single in Figure 5 E).

## References

Anderson, M. J., Yao, A., Laslow, B., Schipani, E., & Lewandoski, M. (2025). HIF1α controls somitogenesis and spine development by regulating levels of intracellular oxygen in the presomitic mesoderm. bioRxiv, 2025–2012.

Aulehla, A., Wehrle, C., Brand-Saberi, B., Kemler, R., Gossler, A., Kanzler, B., & Herrmann, B. G. (2003). Wnt3a plays a major role in the segmentation clock controlling somitogenesis. Developmental Cell, 4(3), 395–406.

Bessho, Y., Sakata, R., Komatsu, S., Shiota, K., Yamada, S., & Kageyama, R. (2001). Dynamic expression and essential functions of Hes7 in somite segmentation. Genes & Development, 15(20), 2642–2647.

Bone, R. A., Bailey, C. S., Wiedermann, G., Ferjentsik, Z., Appleton, P. L., Murray, P. J., Maroto, M., & Dale, J. K. (2014). Spatiotemporal oscillations of Notch1, Dll1 and NICD are coordinated across the mouse PSM. Development, 141(24), 4806–4816.

Boutros, M., Heigwer, F., & Laufer, C. (2015). Microscopy-based high-content screening. Cell, 163(6), 1314–1325.

Brent, A. E., & Tabin, C. J. (2002). Developmental regulation of somite derivatives: Muscle, cartilage and tendon. Current Opinion in Genetics & Development, 12(5), 548–557.

Chal, J., & Pourquié, O. (2009). Patterning and differentiation of the vertebrate spine. The Skeletal System, 53, 41–116.

Christ, B., & Scaal, M. (2008). Formation and differentiation of avian somite derivatives. Somitogenesis, 1–41.

Chu, L.-F., Mamott, D., Ni, Z., Bacher, R., Liu, C., Swanson, S., Kendziorski, C., Stewart, R., & Thomson, J. A. (2019). An in vitro human segmentation clock model derived from embryonic stem cells. Cell Reports, 28(9), 2247–2255.

Delaune, E. A., François, P., Shih, N. P., & Amacher, S. L. (2012). Single-cell-resolution imaging of the impact of notch signaling and mitosis on segmentation clock dynamics. Developmental Cell, 23(5), 995–1005.

Dequéant, M.-L., Glynn, E., Gaudenz, K., Wahl, M., Chen, J., Mushegian, A., & Pourquié, O. (2006). A complex oscillating network of signaling genes underlies the mouse segmentation clock. Science, 314(5805), 1595–1598.

Diaz-Cuadros, M., Wagner, D. E., Budjan, C., Hubaud, A., Tarazona, O. A., Donelly, S., Michaut, A., Al Tanoury, Z., Yoshioka-Kobayashi, K., Niino, Y., et al. (2020). In vitro characterization of the human segmentation clock. Nature, 580(7801), 113–118.

Ho, C., Jutras-Dubé, L., Zhao, M. L., Mönke, G., Kiss, I. Z., François, P., & Aulehla, A. (2024). Nonreciprocal synchronization in embryonic oscillator ensembles. Proceedings of the National Academy of Sciences, 121(36), e2401604121.

Hubaud, A., Regev, I., Mahadevan, L., & Pourquie, O. (2017). Excitable dynamics and yap-dependent mechanical cues drive the segmentation clock. Cell, 171(3), 668–682.

Matsuda, M., Lázaro, J., & Ebisuya, M. (2025). Metabolic activities are selective modulators for individual segmentation clock processes. Nature Communications, 16(1), 845.

Matsuda, M., Yamanaka, Y., Uemura, M., Osawa, M., Saito, M. K., Nagahashi, A., Nishio, M., Guo, L., Ikegawa, S., Sakurai, S., et al. (2020). Recapitulating the human segmentation clock with pluripotent stem cells. Nature, 580(7801), 124–129.

Meijer, H. A., Hetherington, A., Johnson, S. J., Gallagher, R. L., Hussein, I. N., Weng, Y., Rae, J. M., Noordzij, T. E., Kalamara, M., Macartney, T. J., et al. (2025). NOTCH1 S2513 is critical for the regulation of NICD levels impacting the segmentation clock in hiPSC-derived PSM cells and somitoids. Genes & Development, 39(17-18), 1025–1044.

Miao, Y., Djeffal, Y., De Simone, A., Zhu, K., Lee, J. G., Lu, Z., Silberfeld, A., Rao, J., Tarazona, O. A., Mongera, A., et al. (2023). Reconstruction and deconstruction of human somitogenesis in vitro. Nature, 614(7948), 500–508.

Miyazawa, H., Rada, J., Prior, N., Sanchez, P. G. L., Esposito, E., Bunina, D., Girardot, C., Zaugg, J., & Aulehla, A. (2025). A noncanonical role of glycolytic metabolites controlling the timing of mouse embryo segmentation. Science Advances, 11(38), eadz9606.

Palmeirim, I., Henrique, D., Ish-Horowicz, D., & Pourquié, O. (1997). Avian hairy gene expression identifies a molecular clock linked to vertebrate segmentation and somitogenesis. Cell, 91(5), 639–648.

Pourquié, O. (2022). A brief history of the segmentation clock. Developmental Biology, 485, 24–36.

Sanaki-Matsumiya, M., Matsuda, M., Gritti, N., Nakaki, F., Sharpe, J., Trivedi, V., & Ebisuya, M. (2022). Periodic formation of epithelial somites from human pluripotent stem cells. Nature Communications, 13(1), 2325.

Sanchez, P. G. L., Mochulska, V., Mauffette Denis, C., Mönke, G., Tomita, T., Tsuchida-Straeten, N., Petersen, Y., Sonnen, K., François, P., & Aulehla, A. (2022). Arnold tongue entrainment reveals dynamical principles of the embryonic segmentation clock. Elife, 11, e79575.

Sewell, W., Sparrow, D. B., Smith, A. J., Gonzalez, D. M., Rappaport, E. F., Dunwoodie, S. L., & Kusumi, K. (2009). Cyclical expression of the notch/wnt regulator nrarp requires modulation by Dll3 in somitogenesis. Developmental Biology, 329(2), 400–409.

Stringer, C., Wang, T., Michaelos, M., & Pachitariu, M. (2021). Cellpose: A generalist algorithm for cellular segmentation. Nature Methods, 18(1), 100–106.

Takada, S., Stark, K. L., Shea, M. J., Vassileva, G., McMahon, J. A., & McMahon, A. P. (1994). Wnt-3a regulates somite and tailbud formation in the mouse embryo. Genes & Development, 8(2), 174–189.

Takashima, Y., Ohtsuka, T., González, A., Miyachi, H., & Kageyama, R. (2011). Intronic delay is essential for oscillatory expression in the segmentation clock. Proceedings of the National Academy of Sciences, 108(8), 3300–3305.

Winfree, A. T. (1980). The geometry of biological time (Vol. 2). Springer.

Yaman, Y. I., & Ramanathan, S. (2023). Controlling human organoid symmetry breaking reveals signaling gradients drive segmentation clock waves. Cell, 186(3), 513–527.

Yamanaka, Y., Hamidi, S., Yoshioka-Kobayashi, K., Munira, S., Sunadome, K., Zhang, Y., Kurokawa, Y., Ericsson, R., Mieda, A., Thompson, J. L., et al. (2023). Reconstituting human somitogenesis in vitro. Nature, 614(7948), 509–520.

Zanella, F., Lorens, J. B., & Link, W. (2010). High content screening: Seeing is believing. Trends in Biotechnology, 28(5), 237–245.

